# Molecular simulations unravel the molecular principles that mediate selective permeability of carboxysome shell protein

**DOI:** 10.1101/2020.06.14.151241

**Authors:** Matthew Faulkner, István Szabó, Samantha L. Weetman, Francois Sicard, Roland G. Huber, Peter J. Bond, Edina Rosta, Lu-Ning Liu

## Abstract

Bacterial microcompartments (BMCs) are nanoscale proteinaceous organelles that encapsulate enzymes from the cytoplasm using an icosahedral protein shell that resembles viral capsids. Of particular interest are the carboxysomes (CBs), which sequester the CO_2_-fixing enzymes ribulose-1,5-bisphosphate carboxylase/oxygenase (Rubisco) to enhance carbon assimilation. The carboxysome shell serves as a semi-permeable barrier for passage of metabolites in and out of the carboxysome to enhance CO_2_ fixation. How the protein shell directs influx and efflux of molecules in an effective manner has remained elusive. Here we use molecular dynamics and umbrella sampling calculations to determine the free-energy profiles of the metabolic substrates, bicarbonate, CO_2_ and ribulose bisphosphate and the product 3-phosphoglycerate associated with their transition through the major carboxysome shell protein CcmK2. We elucidate the electrostatic charge-based permeability and key amino acid residues of CcmK2 functioning in mediating molecular transit through the central pore. Conformational changes of the loops forming the central pore may also be required for transit of specific metabolites. The importance of these *in-silico* findings is validated experimentally by site-directed mutagenesis of the key CcmK2 residue Serine 39. This study provides insight into the mechanism that mediates molecular transport through the shells of carboxysomes, applicable to other BMCs. It also offers a predictive approach to investigate and manipulate the shell permeability, with the intend of engineering BMC-based metabolic modules for new functions in synthetic biology.

## Introduction

Bacterial microcompartments (BMCs) are proteinaceous organelles that sequester key metabolic pathways in the cytoplasm to enhance metabolic performance^1^. These metabolic modules are ubiquitous in many bacterial phyla and play vital roles in CO_2_ fixation, pathogenesis, and microbial ecology^2–4^. BMCs are composed of catalytic enzymes encapsulated by a single-layer protein shell that structurally resembles virus capsids. The shell is self-assembled by hundreds of proteins in the forms of hexamers (BMC-H), pentamers (BMC-P) and trimers (BMC-T). BMC-H and BMC-T proteins are predominant in the shell facets, whereas BMC-P proteins occupy the vertices of the polyhedral shell^5^.

The most well-characterized BMC is the carboxysome, which acts as the key CO_2_-fixing machinery in all cyanobacteria and many chemoautotrophs^6–8^. The primary carboxylating enzymes, ribulose-1,5-bisphosphate carboxylase/oxygenase (Rubisco), are encased in the carboxysome^9^. Based on their protein composition and phylogeny of Rubisco enzymes, carboxysomes can be categorized into two types, α-carboxysomes and β-carboxysomes^10^. The β-carboxysome from freshwater cyanobacteria exhibits an icosahedral geometry with a diameter of ~150 nm (Fig. 1A)^11,12^.

**Fig. 1.**
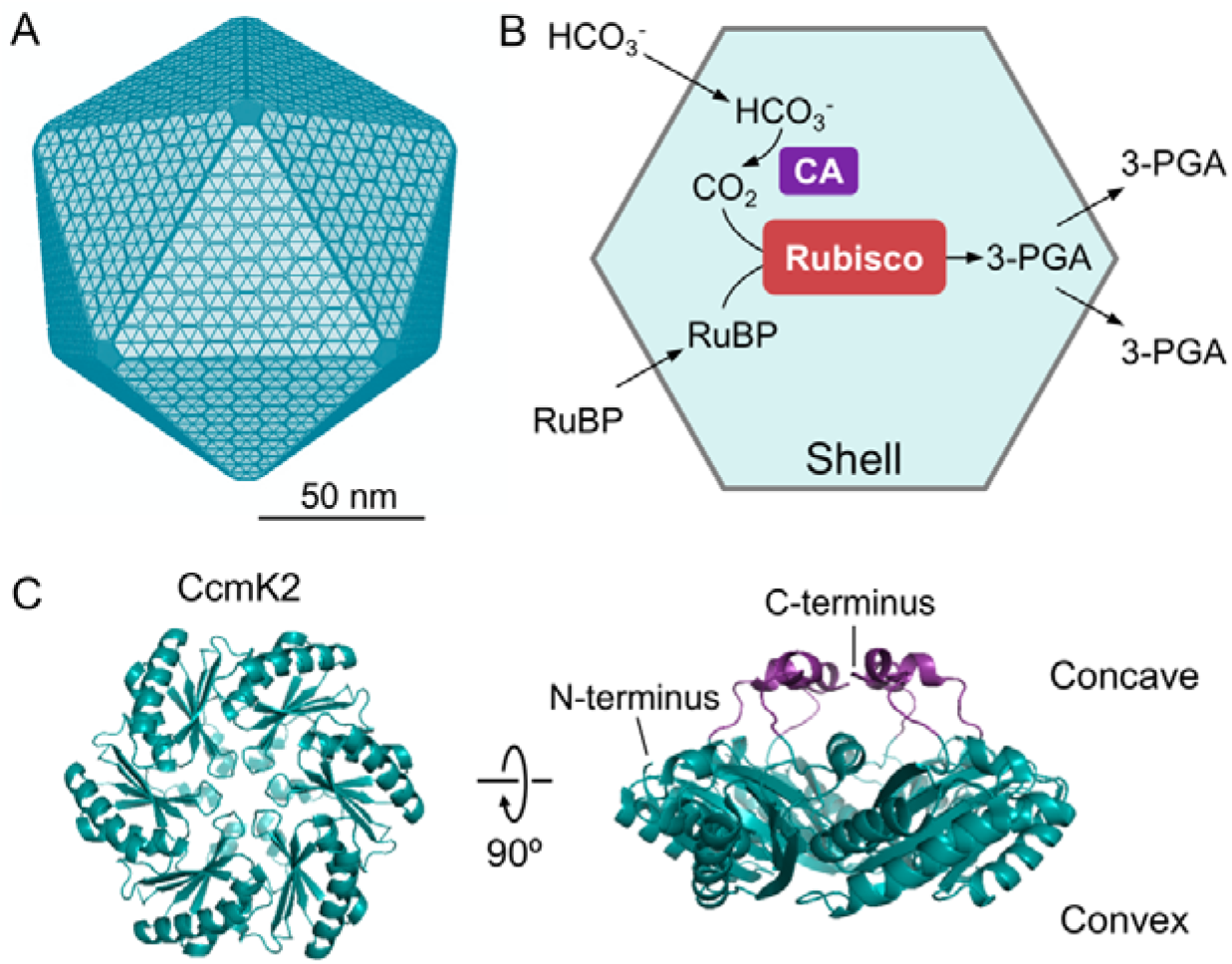
Carboxysome structure, metabolism and building components. **A**, A model of the proposed polyhedral architecture of the carboxysome. The β-carboxysome from Syn7942 is ~150 nm in diameter, comprising an icosahedral shell formed by hexameric and pentametric proteins. **B**, The metabolic pathways of carbon fixation in the carboxysome. The carboxysome shell serves as a physical barrier for controlling the flux of specific metabolites in and out of the carboxysome. The shell permits passage of cytosolic bicarbonate (HCO_3_^−^) and ribulose-1,5-bisphosphate (RuBP) into the carboxysome. Then carbonic anhydrase (CA) in the carboxysome lumen dehydrates HCO_3_^−^ to CO_2_ and provides high levels of CO_2_ close to Rubisco. Rubisco catalyzes the carboxylation of RuBP by adding CO_2_ to generate 3-phosphoglycerate (3-PGA), which is transported across the shell and is metabolized via the Calvin–Benson–Bassham cycle. **C**, Structural representation of the major carboxysome shell building component, CcmK2 (PDB: 2A1B). The CcmK2 complex presents a 6-fold symmetry, involving a central pore that mediates metabolite flow. CcmK2 has concave and convex faces; the C-terminal tails of CcmK2, highlighted in purple, are located at the concave side.

Rubisco catalyzes the reaction between ribulose-1,5-bisphosphate (RuBP) and CO_2_ to generate two molecules of 3-phosphoglycerate (3-PGA). Meanwhile, Rubisco can also carry out oxygenation by accepting O_2_ as a substrate instead of CO_2_ through a competing, unproductive reaction called photorespiration. Its restricted capability in discriminating between CO_2_ and O_2_ causes the catalytical inefficiency of Rubisco^13^. The carboxysome provides a microenvironment that can sequester and concentrate Rubisco enzymes from the cytoplasm (Fig. 1B); carbonic anhydrase (CA) is co-encapsulated with Rubisco within the carboxysome lumen and dehydrates HCO_3_^−^ to CO_2_ and thereby, supplies a high concentration of CO_2_ in the vicinity of Rubisco. The shell of the carboxysome was also speculated to act as a diffusion barrier to HCO_3_^−^/CO_2_ efflux and O_2_ influx, thereby promoting Rubisco carboxylation^14^. The shell should also permit transit of the natural substrate RuBP and product 3-PGA. Overall, the highly-defined structure and protein organization of carboxysomes enhances the efficiency of CO_2_ fixation and reduces the wasteful side reaction of photorespiration, allowing cyanobacteria to contribute up to 25% of global carbon fixation on Earth^15^. As such, there is a growing interest in repurposing carboxysomes in heterologous systems, such as *Escherichia coli* (*E. coli*) and crop plants, to profoundly boost photosynthetic carbon fixation and productivity^16–20^.

All shell proteins bear a central pore surrounded by the loops of BMC-H, BMC-P and BMC-T polypeptides. These pores are presumed to provide portals for metabolite transport in and out of the carboxysome, given that shell proteins are closely packed in flat facets on the basis of observations from crystallography^21,22^, electron microscopy^11,23,24^ and atomic force microscopy^11,25,26^. To date, how the transition of substrates through the carboxysome shell is mediated remains enigmatic. Here, we use molecular dynamics (MD) via umbrella sampling (US) free energy calculations to explore the selective permeability of the major shell protein CcmK2 (BMC-H) in β-carboxysomes to specific substrates and products related to the reaction catalyzed by Rubisco. This study provides computational evidence for elucidating the mechanisms that mediate metabolite traffic through the carboxysome shell.

## Results

### Structural flexibility of CcmK2

CcmK2 in β-carboxysomes from *Synechocystis sp*. PCC 6803 (Syn6803) was the first BMC protein that has been structurally determined^22^. The protein subunits of CcmK2 are arranged in hexameric units with a six-fold symmetry (Fig. 1C). The central pore is ~ 7 Å in diameter, surrounded by six copies of the backbone amide nitrogen atom of Serine 39 (Ser39). The predominant protein surface of CcmK2 is negatively charged, whereas the central pore of CcmK2 has a strong positive electrostatic potential, ascribed to the positively charged residues Arginine 11 (Arg11) and Lysine 36 (Lys36) that are 10 Å away from the narrowest point of the pore in each subunit and solvent exposed to the lumen and cytoplasm, respectively^22^ (Supplementary Fig. 1 and 2).

The CcmK2 C-terminal tail contains a 5-residue helix on the concave side and has been deduced to play roles in BMC domain assembly and protein-protein interaction^22^ (Fig. 2A and 2B). We performed MD simulations of the full-length CcmK2 from Syn6803 (CcmK2-full, PDB: 2A1B)^22^, the C-terminal truncated CcmK2 from Syn6803 (CcmK2-ΔC, PDB: 3CIM)^27^, and the full-length CcmK2 from *Synechococcus elongatus* PCC 7942 (Syn7942) (PDB: 4OX7) in explicit water for 200 ns. The results revealed that the C-terminal tails of CcmK2 exhibited structural flexibility during simulations (Fig. 2C). These C-terminal residues have a lower resolution and display a higher degree of flexibility than the core residues in the crystal structures, resulting in the disorder of crystal forms^27,28^. A similar fashion of protein contacts has been illustrated in virus capsid assembly: interactions through flexible termini act as conformational switches that mediate virus capsid formation^29,30^.

**Fig. 2.**
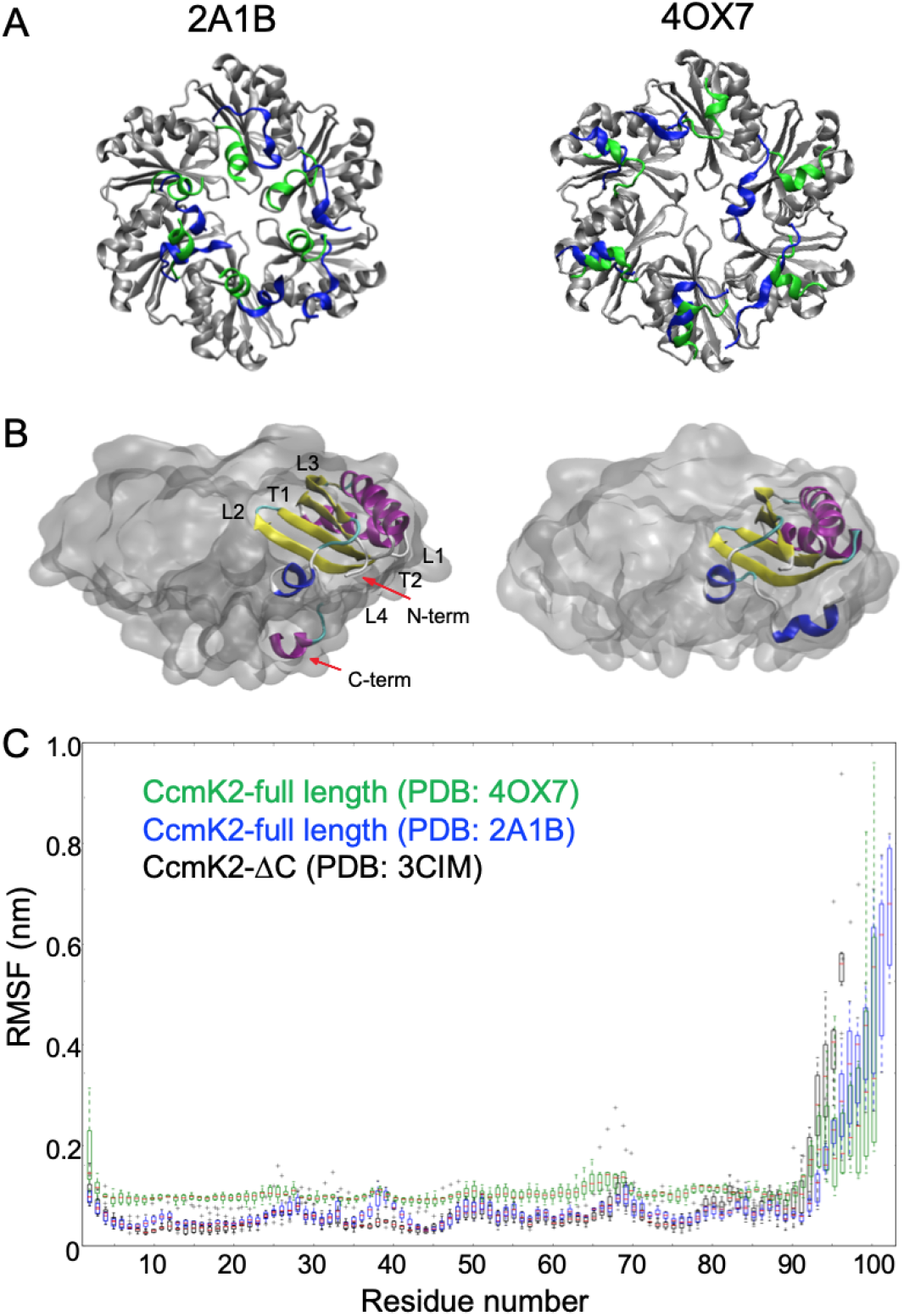
Structural flexibility of the C-terminal tails of CcmK2. **A**, C-terminal orientation of CcmK2-full. The initial orientation of the C-termini is shown in green and the final state of the C-termini after 200 ns of simulation time is shown in blue. **B**, Visualization of an individual chain (cartoon representation) in the hexamer (quick-surf representation) with the termini, loop and turn regions labeled. **C**, RMSF profile of the truncated CcmK2 (PDB: 3CIM) and full-length CcmK2 (PDB: 2A1B and 4OX7) over the last 150 ns of the simulation period, indicating a flexible C-terminus of CcmK2.

We conducted root mean square fluctuation (RMSF) analysis to measure the per residue fluctuation around their average positions. High values of RMSFs are seen at the C-termini (Fig. 2C), as a consequence of the high C-terminal flexibility during the course of simulations. Other regions of increased RMSF peaks correspond to regions of high flexibility such as loops and turns in CcmK2. By contrast, no significant difference in the RMSF profiles of the core regions of CcmK2 was recorded, indicative of the stable conformation of the core. To diminish the system complexity, we chose the experimentally obtained CcmK2-ΔC structure (PDB: 3CIM)^27^, with a high resolution of 1.3 Å, in the following simulations to explore the molecular transport through the central pore, instead of creating *in silico* a C-terminal deletion structure based on the full-length CcmK2. Since the MD simulation lengths cannot sample the position of the flexible termini exhaustively to find a stable conformation relative to the whole protein, using the CcmK2-ΔC structure eliminates this uncertainty.

### Molecular transport of the pore is mediated by electrostatic charge

The pore of CcmK2 has been proposed to be the route by which metabolites pass through the shell in either direction^22^. We conducted US simulations on CcmK2-ΔC (PDB: 3CIM) in the solvent containing HCO_3_^−^, 3-PGA, O_2_ and CO_2_, respectively. We determined the free-energy (ΔG) profiles associated with the transition of each metabolite initiating simulations in both directions, along the axis (*z*) relative to the center of mass of the CcmK2 hexamer (*x*, *y*, *z*) = (0, 0, 0) (Fig. 3A). To assess the US convergence, we divided each window into 5 equal-length trajectories to calculate the error bars, which did not display significant systematic changes (Supplementary Fig. 4). We also analyzed the autocorrelation functions at 3 different US windows (Supplementary Fig. 5) and obtained around 2 ns for the autocorrelation times, shorter than the US simulation lengths.

**Fig. 3.**
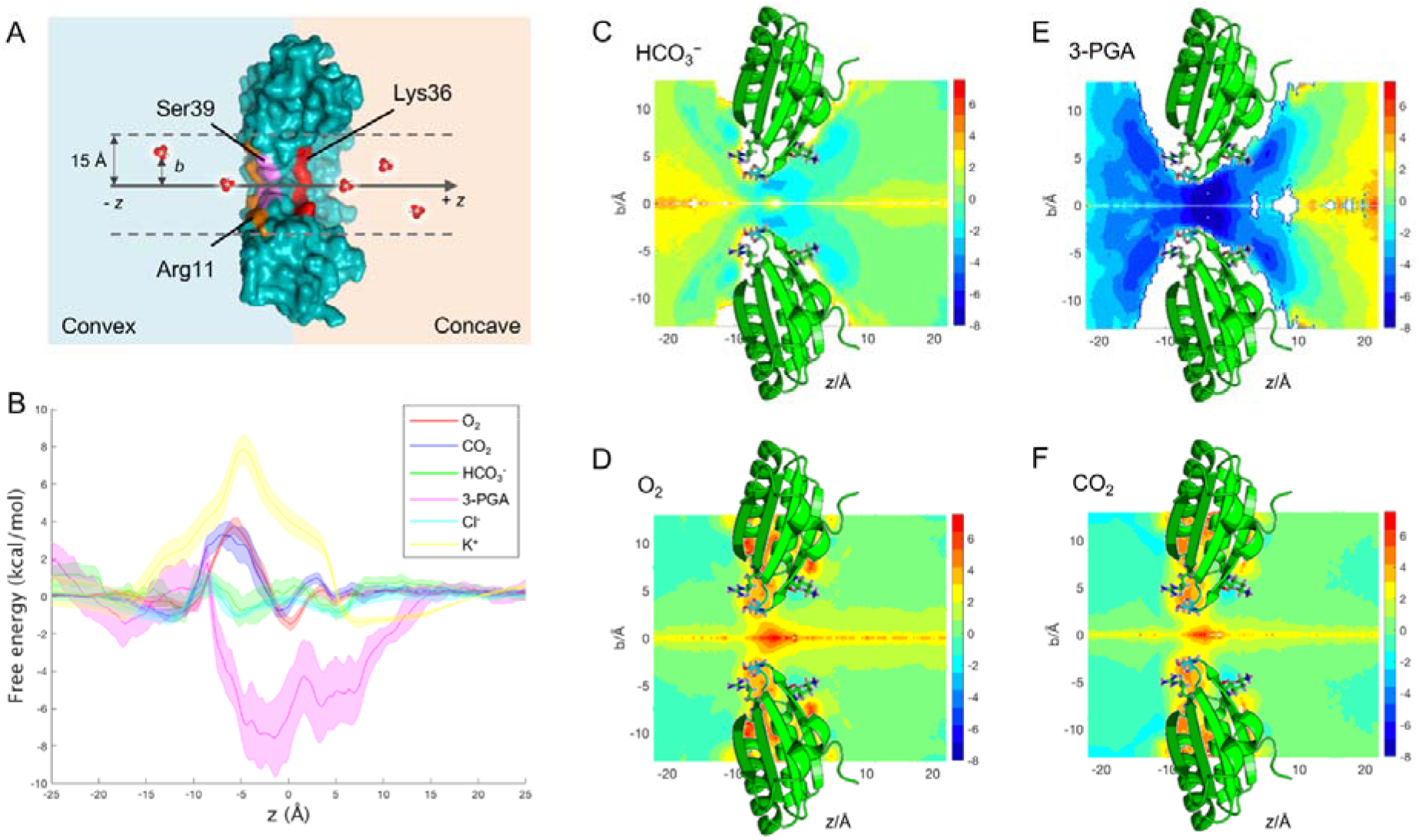
Selective permeability of the CcmK2-ΔC pore towards key metabolites and products related to carbon fixation. **A**, A cross-section through a schematic representation of the US environment. The *z* axis shown is relative to (0,0,0) at the center of mass of CcmK2. The full range sampled was from −25 to +25 Å along *z*. Positive values of *z* refer to the concave side of CcmK2, whereas negative values of *z* refer to the convex surface. **B**, Free-energy profiles of key metabolite molecules HCO_3_^−^, 3-PGA, O_2_ and CO_2_ along *z* axis, and of K^+^ cations and Cl^−^ anions from no-metabolite control environments. C-F, 2D free-energy landscapes of HCO_3_^−^, O_2_, 3-PGA and CO_2_ transport, respectively. A cartoon representation of CcmK2 is shown in green with the side chains of the three key residues Ser39, Lys36 and Arg11 shown as sticks.

As shown in Fig. 3B, the local free energy differences are calculated relative to bulk solvent (at *z* = −25 Å, +25 Å). It is evident that the four metabolites experience three shared local free-energy minima/maxima, representing their specific interactions with the key amino acid residues (Fig. 3). O_2_ and CO_2_ have the same local free-energy minima localized around the side chains of Arg11 (*z* = ~−11 Å) and Lys36 (*z* = ~5 Å) as well as a free energy barrier, ΔG^‡^ of 3.8 kcal⋅mol^−1^ and 3.5 kcal⋅mol^−1^ respectively, in proximity to Ser39 (*z* = ~−6 Å) (Fig. 3B). This indicates that both the CO_2_ and O_2_ has to pass a small free energy barrier to transition through the pore from one side of Ser39 to the other.

The free-energy minima of HCO_3_^−^ are around the side chains of residues near the center of the pore, which is formed by the K-I-G-S motif^31^; Ser39 where *z* is ~−5 Å, Arg11 where *z* is ~−12 Å, and Lys36 where *z* is ~6 Å. The lowest of the energy wells occurs at the Ser39 bottle neck, with a ΔG of ~−0.6 kcal⋅mol^−1^. It is the same for 3-PGA except that there is no minimum at Arg11. These free-energy minima correlate with the binding pockets of CcmK2 for metabolites in 3D space (Fig. 3C-3F), indicating the importance of residues Ser39, Lys36 and Arg11 in mediating metabolite flux through the pore. Consistently, recent studies on 1,2-propanediol utilization (1,2-PDU) microcompartments have also verified the role of the equivalent Ser40 of the shell hexamer PduA in channeling metabolite traffic^32,33^. Given that the Ser39, Lys36 and Arg11 residues are conserved among orthologous BMC-H members as demonstrated by protein sequence alignment (Supplementary Fig. 3)^22,27^, our results may suggest a general principle underlying the selective permeability of BMC shell proteins and the conservation of functional domains of BMC shell components. Collectively, our results reveal the charge-based molecule transport through the electropositive CcmK2 pore.

Additionally, we calculated the free energy of K^+^ and Cl^−^ using metabolite-free US simulations. The cation K^+^ displays a maximum between *z* = ~−16 to 5, with the highest ΔG ~7.6 kcal⋅mol^−1^ occurring around Ser39 *z* = ~−4. This suggests that it is unfavorable for cations to transition through the CcmK2 pore. In contrast, the small anion Cl^−^ has the smoothest energy profile and is most similar to that of HCO_3_^−^. Overall, it experiences much smaller barrier heights for a favorable transition (Fig. 3B). The most favorable free energy occurs when 3-PGA transits around Ser39 with a ΔG of ~−7.5 kcal⋅mol^−1^(Fig. 3B). This could be rationalized by the tendency for hydrogen bonds to be formed between the carboxyl group of 3-PGA and the hydroxyl group of Ser39. The local minimum of 3-PGA on the convex side is higher than around Lys36 on the concave face of CcmK2. In contrast, the local minimum of HCO_3_^−^ around Arg11 (*z* = −12 Å) on the convex face of CcmK2 is approximately equal to the minimum around Lys36 (*z* = 6 Å) on the concave side. There is also a shared local maximum centered at *z* = 0–2 Å between the minima around Lys36 and the minima at Ser39.

These results suggest that energetically favorable interactions around and in the center of the pore, increase the likelihood of transit and could drive transit of HCO_3_^−^ and 3-PGA in both directions. It is possible that 3-PGA preferentially transits from the convex to the concave face. The energy profile suggests more favorable interactions occur on the concave side of Ser39 (Fig. 3B). Whereas HCO_3_^−^ US doesn’t indicate a preference, it is likely in the cellular environment HCO_3_^−^ would preferentially transit from the concave to the convex face with a concentration gradient. This is in agreement with the shell protein orientation characterized in the intact BMC shells^34^ and the PduA nanotubes^35^, but distinct from the proposed orientation in recent simulations of the α-carboxysome shell hexamer CsoS1A^36^. The pore, and the binding pockets at the key residues on either side, are accessible to water. Solvent effects, from the likes of salinity, merit further investigation (Fig. 4, Supplementary Fig. 2).

**Fig. 4.**
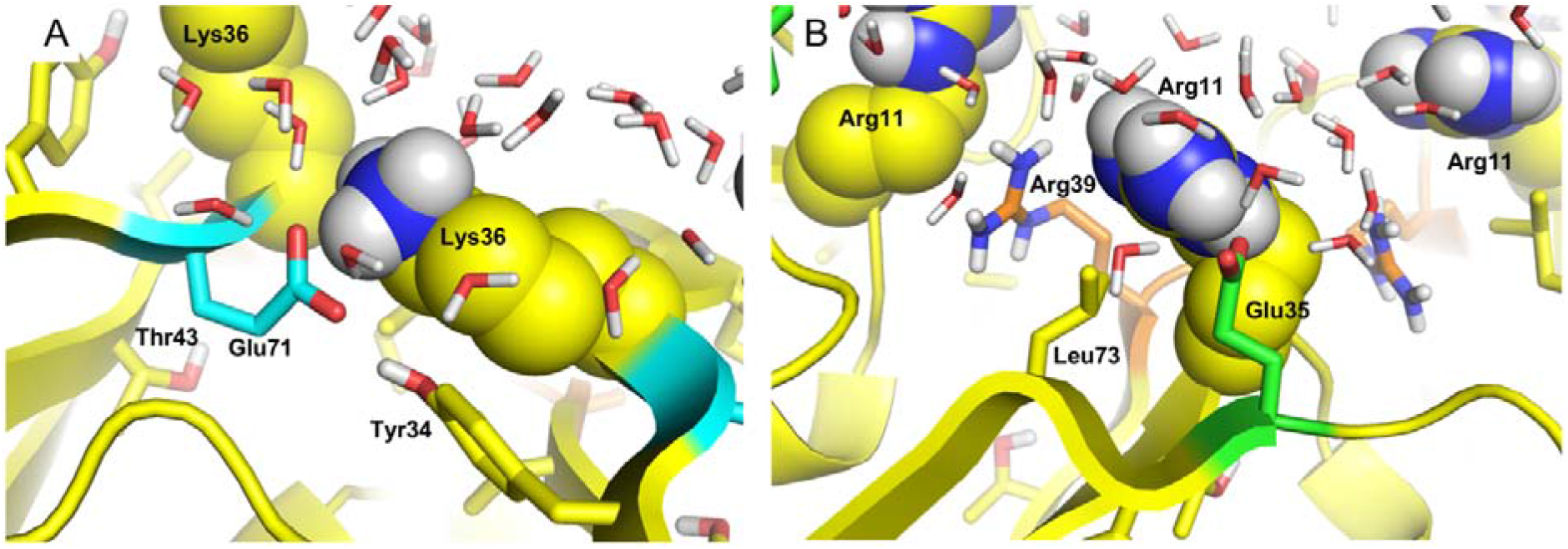
Interactions formed by key charged residues. Lys36 (left) and Arg11 (right) shown as VDW spheres at the concave and convex sides of the pore respectively. Interacting conserved Glu71 (turquoise) and Glu35 (green) residues, as well as nearby residues within 4 Å (Tyr34 and Arg41, Leu73) and water molecules are shown as sticks highlighting solvent accessibility.

### Conformational change of the central pore during metabolite transit

To examine whether potential conformational changes of the CcmK2 pore are associated with transit of metabolites, we measured the pore diameter in the presence and absence of contacts between individual metabolites and the residues Arg11, Ser39 and Lys36 during the time course of simulations (Fig. 5), using HOLE^37^ to analyze 2500 equidistant frames of a 50 ns MD trajectory independently (see Materials and Methods). The pore size of CcmK2-ΔC in the crystal structure (PDB: 3CIM) was 2.55 Å determined using HOLE, and it varied between 1 and 6 Å during the course of simulations (Fig. 5). There were no significant differences found in the pore diameter in the absence and presence of contacts between CO_2_ or O_2_ and the three residues. It is worth noting that the sample size of contacts between these metabolites and the select residues is small, likely because the substrates with relatively small size have more space to move freely in the pore area without making contacts. On the contrary, a significant difference in the pore size was discerned for HCO_3_^−^ between no contacts and contacts with Lys36 (Fig. 5, Supplementary Table 2). This reveals that transit of HCO_3_^−^ may first require binding to Lys36 on the concave side of the protein to open up the pore. Significant changes in the pore diameter were also detected in the presence and absence of contacts between 3-PGA and all three of the selected residues of interest, indicating that the passage of 3-PGA through the pore may require a conformational change to increase the pore diameter (Fig. 5).

**Fig. 5.**
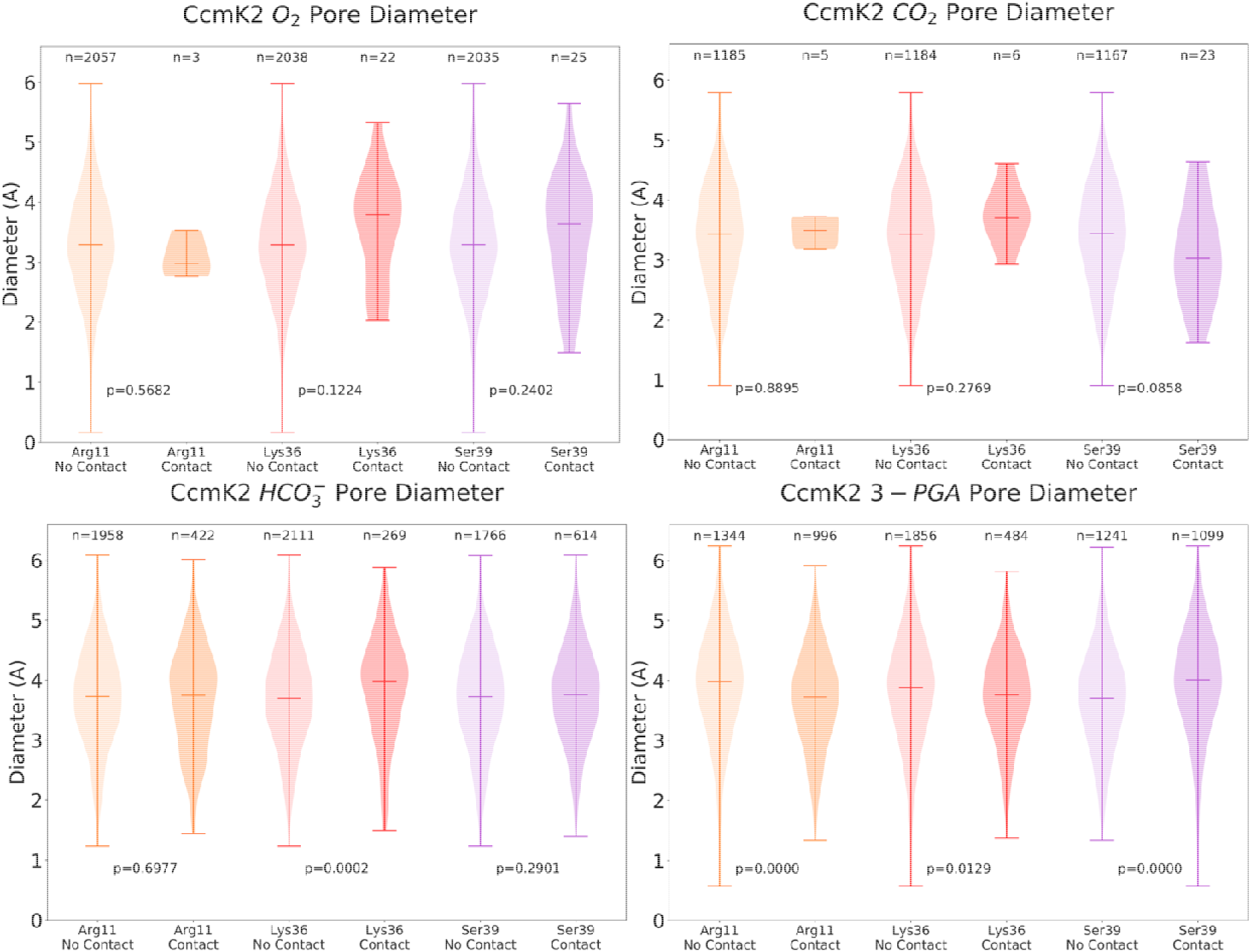
Pore diameter of CcmK2-ΔC in the presence and absence of contacts with metabolites during simulations. Violin plots of the pore diameters of CcmK2 determined by HOLE from the full MDS trajectories for each metabolite, depicting the conformational dynamics of the pore during metabolite transition. The two categories “contact” and “no contact” refer to the presence or absence, respectively, of the metabolite within the cutoff distance of 3.5 Å and angle of 30° of the reside. *p* values were calculated using the Wilcoxon rank-sum test (Supplementary Table 2). These diameter measurements were made at the region corresponding to the minimum constriction along the pore axis *z*.

### Permeability of large metabolite molecules through the CcmK2 pore

Our results imply that the large molecule 3-PGA experiences favorable interactions (Fig. 3B) and may induce a conformational change of the CcmK2 pore during transit (Fig. 5). Likewise, RuBP also has a large size (~3-4 Å wide, ~9.4 Å long) with respect to the pore diameter (Fig. 6A). To estimate their interactions with the CcmK2 pore, we conducted molecular docking simulations of 3-PGA and RuBP with CcmK2-ΔC structure using SwissDock^38^ and Glide extra precision (XP) docking^39^. Snapshots of the most stable binding poses for RuBP (Fig. 6B) and 3-PGA (Fig. 6C) from both the concave and convex faces illustrate the negative relative free energy changes experienced by RuBP and 3-PGA when in the final docking pose in the CcmK2 pore: Δ*G* is −9.96 kcal·mol^−1^ for RuBP (Fig. 6D) and −9.98 kcal·mol^−1^ for 3-PGA (Fig. 6E). The FullFitness score for RuBP (−3383 kcal·mol^−1^) in this pose is similar to 3-PGA (−3275 kcal·mol^−1^) (Supplementary Fig. 6). These results verify a highly stable binding pocket for 3-PGA and RuBP in the narrowest point of the CcmK2 pore when the phosphate groups of these metabolites interact with the Ser39 hydroxyl groups. All three of the phosphate oxygen groups of both RuBP and 3-PGA can be stabilized by hydrogen bonding to Ser39. This binding pocket also contributes to a local free-energy minimum for 3-PGA, as observed in US calculations (*z =* ~−5 Å) (Fig. 3). This emphasizes a good correlation between our molecular docking and US simulations results, despite the known lower reliability of docking simulations compared to full MD simulations. These binding poses of 3-PGA are directly compared between US and docking in Fig. 7. Interestingly, despite the size difference between 3-PGA and RuBP, the free energy detected when docking RuBP and 3-PGA to CcmK2 is relatively similar. This might suggest that there are relatively strong interactions between the binding pocket and RuBP or 3-PGA. This binding pocket is alike for both molecules, but accessibility is equally challenging due to the small diameter of the pore and large sizes of the metabolites. As the binding poses observed are solvent-accessible and make use of the charged phosphate groups of the metabolites bound in the polar center of the pore, we do not expect significant changes in their protonation states upon binding. Nevertheless, a significant change in pH that affects the protonation states of these metabolites might result in the changes in binding energies and poses.

**Fig. 6.**
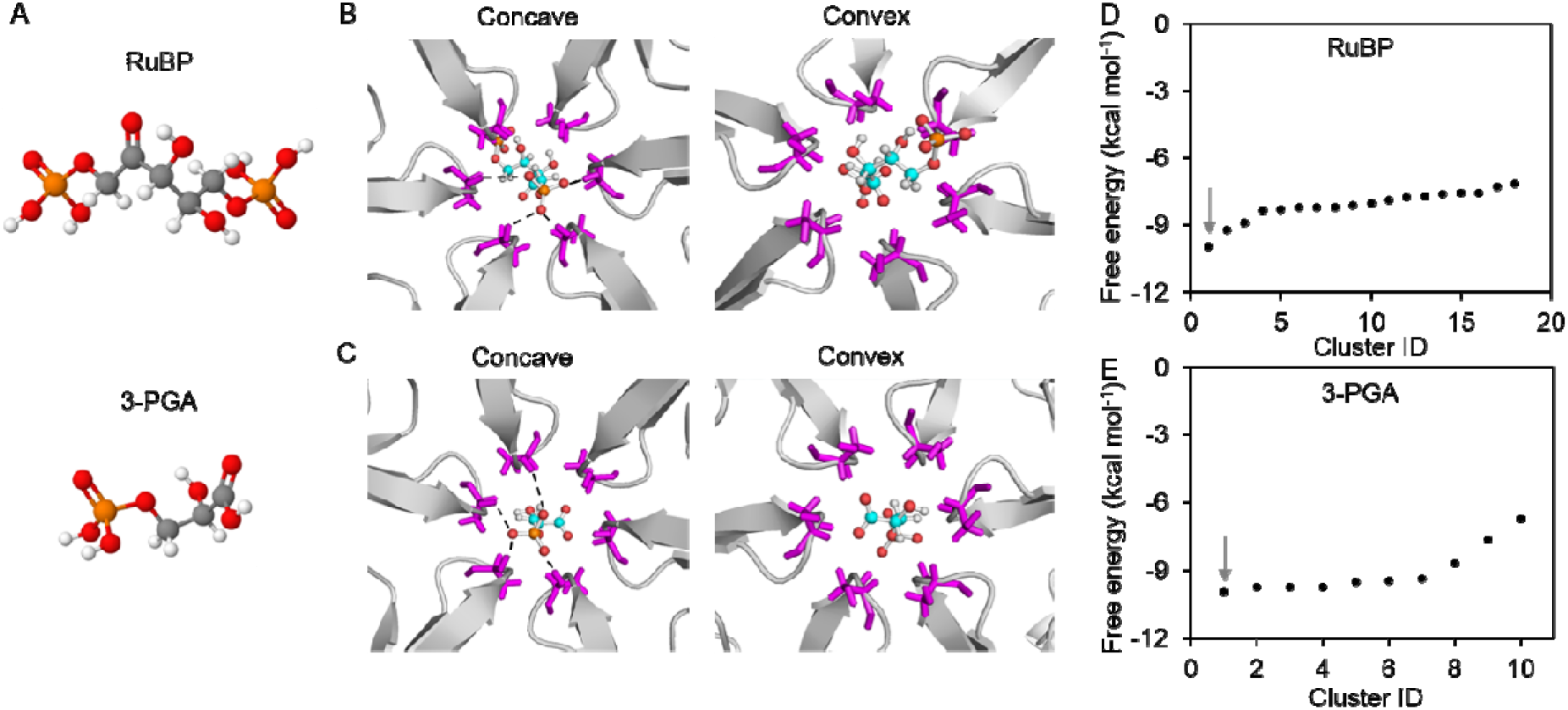
Interactions of large metabolites RuBP and 3-PGA with CcmK2-ΔC. **A**, Molecular structures of RuBP and 3-PGA. **B**, Interactions of RuBP with the Ser39 residues (purple) displayed from concave and convex sides, hydrogen bonding between the phosphate oxygens and Ser39 side chain indicated by dotted lines. **C**, Interactions of 3-PGA with the Ser39 residues (purple) displayed from concave and convex sides. **D**, Free energy scores of clustered RuBP binding positions. The interaction site shown in **B** bears the minimal free energy in Cluster ID 1, as indicated by the arrow. **E**, Free energy scores of clustered 3-PGA binding positions. The interaction site shown in **C** has the minimal free energy in Cluster ID 1, as indicated by the arrow. The results suggest that the most stable interacting sites with RuBP and 3-PGA are both on the convex side of CcmK2.

**Fig. 7.**
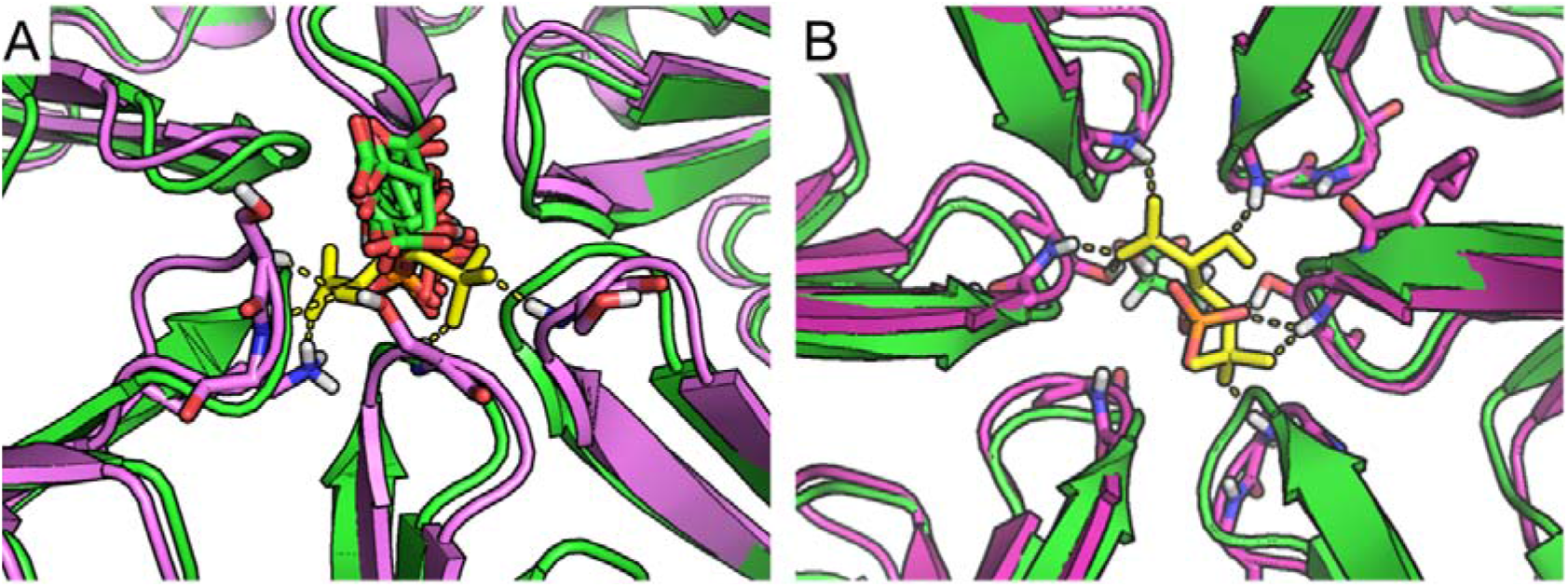
Comparison of docking and US MD poses of 3-PGA. **A**, View from the concave side, docking poses (green cartoon for protein and sticks for 3-PGA) compared with 3-PGA bound structures at the free energy minimum from US MD simulations (3-PGA: yellow sticks, protein: purple cartoon). Note that more extended hydrogen bonds can form due to small rotations of the loops at the center of the pore, allowing interactions of 3-PGA with Ser backbones, Lys and Ser sidechains (shown as sticks). **B**, Comparison between top docked pose (green cartoon for protein and sticks) and selected MD snapshot from the free energy minimum (purple cartoon and yellow sticks).

### Effects of pore mutation on *in vivo* physiology

Our simulation results suggest that the Ser39 residue in the CcmK2 is a key residue in the charged-based mechanism that mediates the flux of metabolites into and out of the carboxysome. To corroborate the *in-silico* findings, we investigated the significance of Ser39 on carboxysome activities and cell physiology by constructing a CcmK2-S39A point mutation. The carboxysome of Syn6803 in addition to CcmK2 contains another major shell protein CcmK1^21^, whereas in Syn7942, CcmK2 proteins act as the only major shell components on the carboxysome shell, as shown by a previous proteomic study^40^. The CcmK2 proteins of Syn7942 and Syn6803 present a high sequence similarity and the Ser39 residues are highly conserved (Supplementary Fig. 3). To prevent the complementary effects of CcmK1, whose role is still unclear, we generated the CcmK2-S39A mutant in Syn7942 and examined cell growth and carbon fixation activities of the Syn7942 mutants including knockouts of the other shell proteins.

We compared the growth of CcmK2-S39A in three conditions: air, NaHCO_3_ supplemented, and 4% CO_2_. The CcmK2-S39A strain could grow at 4% CO_2_, with a similar growth rate with WT (Fig. 8A). By contrast, the growth of CcmK2-S39A was greatly impeded under the air and HCO_3_^−^ supplemented conditions, compared with those of WT under the same conditions (Fig. 8B). This raises the possibility that the S39A mutation may preclude transport of bicarbonate into carboxysomes, or prevents other functions of the pore, such as luminal CO_2_ retention. We also determined that the distinct carbon-fixation activities of the WT and CcmK2-S39A cells grown under 4% CO_2_ (Fig. 8C). CcmK2-S39A has a lower apparent *V*_*max*_ (10.0 ± 1.5 μmol·min^−1^·OD^−1^, *n* = 4) and a higher *K*_*m*(RuBP)_ (1.7 ± 0.6 mM, *n* = 4) compared with the WT (*V*_*max*_ = 28.3 ± 2.0 μmol·min^−^ 1·OD^−1^, *K*_*m*(RuBP)_ = 0.7 ± 0.4 mM, *n* = 4), revealing the affected Rubisco activity caused by the pore mutation. The increased RuBP binding affinity *K*_*m*(RuBP)_ might act as a physiological adaption to compensate for a decreased propensity of RuBP transit into the carboxysome, especially at a low concentration of RuBP.

**Fig. 8.**
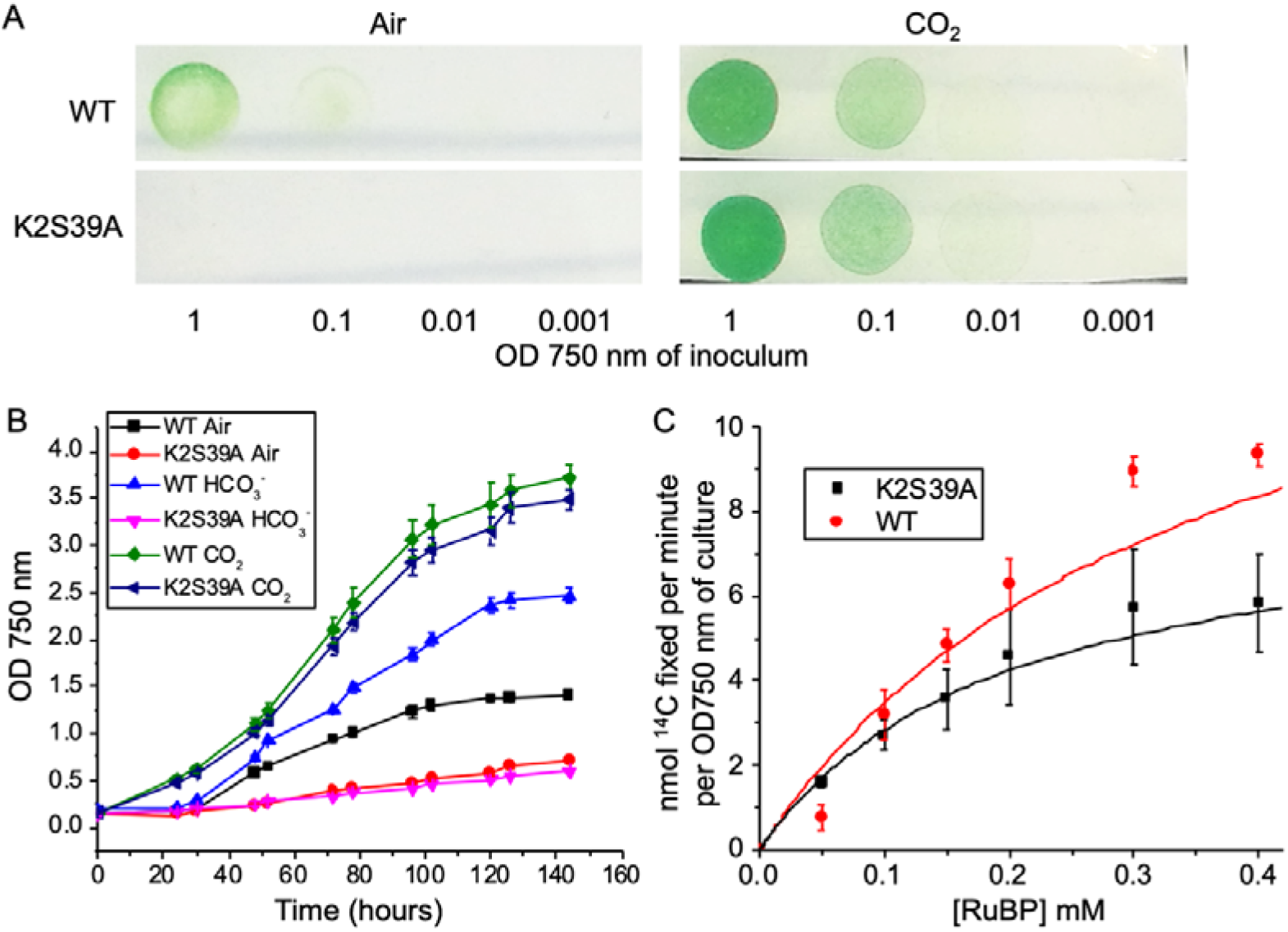
Physiology of the Syn7942 WT and CcmK2-S39A mutant. **A**, Growth of WT and CcmK2-S39A strains under air or 4% CO_2_. A series of dilutions from liquid cultures with OD750 of 1 were plated on BG11 plates and imaged after 48 hours. **B**, Growth curves of WT and CcmK2-S39A cells in air, 4% CO_2_, or with daily 150 mM HCO_3_^−^ supplements. Growth of three biological repeats (*n* = 3) was recorded. CcmK2-S39A requires high CO_2_ to survive, grow slowly in air, and with HCO_3_^−^ supplements. **C**, Carbon fixation activities of WT and CcmK2-S39A (OD750 = 1) as a function of RuBP dosage, fitted with Michaelis-Menten curves (*n* = 4). CcmK2-S39A has a lower *V*_*max*_ (10.0 ± 1.5 μmol·min^−1^·OD^−1^) and a higher *K*_*m*_ (1.7 ± 0.6 mM) compared with WT (*V*_*max*_ = 28.3 ± 2.0 μmol·min^−1^·OD^−1^, *K*_*m*_ = 0.7 ± 0.4 mM).

We further generated the CcmK2-S39A mutation in Δ*ccmK3*, Δ*ccmK4*, Δ*ccmK3K4* and Δ*ccmP* Syn7942 strains, respectively. It is evident that CcmK2-S39A had a notable influence on the growth of all these strains in the air and HCO_3_^−^ supplemented conditions and resulted in a detectable reduction in carbon-fixation activity (Supplementary Fig. 7). The effects of point mutations of CcmK2 Arg11 and Lys36 residues on shell permeability merit further investigation.

## Discussion

Selective permeability of the carboxysome shell allows the generation of a local microenvironment with an optimized luminal concentration of metabolites, crucial for carbon assimilation of this specialized organelle. In particular, the carboxysome shell was proposed to facilitate the diffusion of HCO_3_^−^ and probably provides a barrier to O_2_ and reduces CO_2_ leakage into the cytosol^41^, and thus, plays roles in minimizing the unproductive oxygenation and improving the carboxylation of Rubisco encased by the carboxysome shell. The shell is also permeable to protons, leading to similar pH conditions in the carboxysome compartment lumen and the cytoplasm^42^. Here, we conducted a systematic characterization of the principles underlying the permeability of the major carboxysome shell protein CcmK2^12^, using an array of computational approaches. Our study suggests that the mechanism of metabolite transit through the CcmK2 pore represents a combination of electrostatic charge-based transition and the conformational change of the pore. Our simulations results, combined with the recent findings which have demonstrated that the concave surfaces of shell proteins in the uniformly-oriented shell^25^ face out of the BMC^34,35^, allow us to propose a model for the metabolite flux across the carboxysome shell (Fig. 9).

**Fig. 9.**
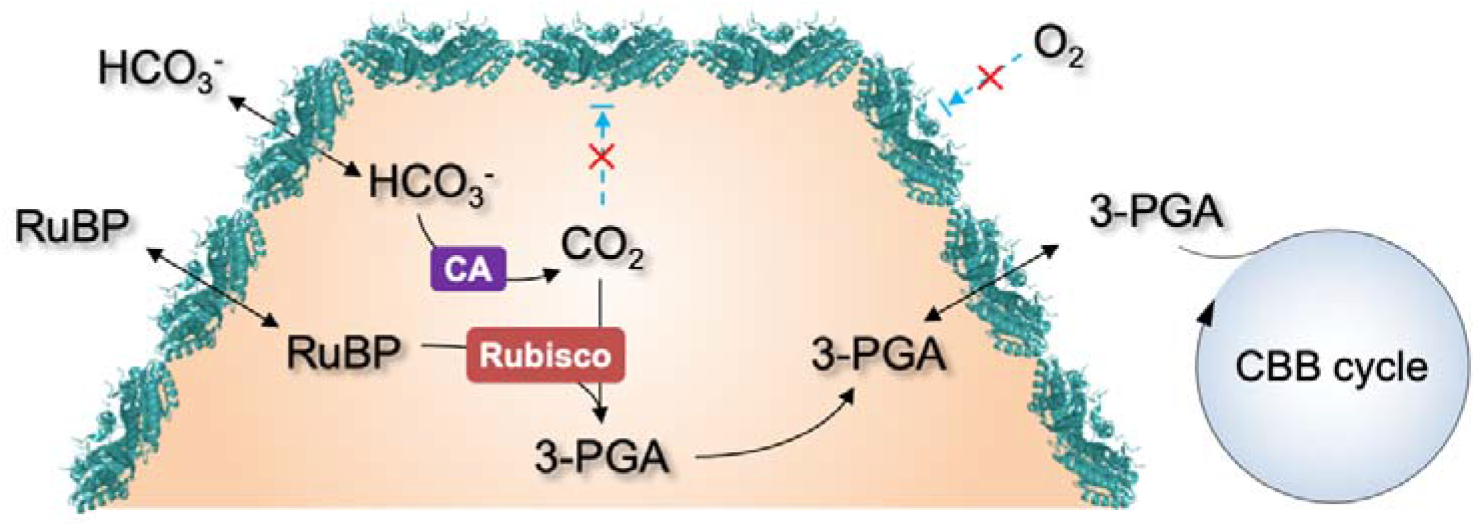
Schematic model representing the selective permeability of the carboxysome shell to confine metabolite flux for driving the CBB cycle. CcmK2 hexamers are the major components in the icosahedral carboxysome shell, with the concave side facing outwards to the cytoplasm and the convex side facing inwards to the lumen. Based on the positive electrostatic charge of the central pore, CcmK2 acts as a tunnel for HCO_3_^−^ influx and a barrier to O_2_ and CO_2_, precluding O_2_ influx and leakage of CO_2_ from the carboxysome lumen to the cytoplasm. Larger molecules RuBP and 3-PGA can likely pass through CcmK2 but require a conformational change in the CcmK2 pore involving a flip of the Ser39 side chain.

We show that the selective permeability of CcmK2 is dependent on the electrostatic charges of molecules. It is energetically favorable for the negatively charged HCO_3_^−^ to move through the pore, but unfavorable for CO_2_ and O_2_ to pass the pore, akin to the experimental findings^41,43^. This is of physiological significance for improving carbon fixation. Bicarbonate represents the dominant fraction of carbon dioxide in the aqueous cyanobacteria-growing environment and the cytoplasm. It is pumped across the cytoplasmic membranes of cyanobacteria into the cytoplasm by HCO_3_^−^ transporters, resulting in a great accumulation of bicarbonate within the cell to a level up to 1000-fold higher than exogenous bicarbonate^44^. The cytosolic HCO_3_^−^ then passes across the shell and is converted to CO_2_ by CA that is co-encapsulated with Rubisco within the carboxysome (Fig. 9), leading to elevated levels of CO_2_ close to Rubisco to favor the carboxylation activities of Rubisco. Although free-energy simulations could not define explicitly the preferable direction of HCO_3_^−^ transit through the pore (Fig. 3), the HCO_3_^−^ gradient across the shell (a high level of HCO_3_^−^ outside the shell and a falloff of the HCO_3_^−^ level within the compartment in the presence of CA) may play the decisive role in driving the flow of HCO_3_^−^ into the carboxysome. In addition, both transits of CO_2_ and O_2_ through the pore were revealed to be energetically unfavorable (Fig. 3). This suggests that the shell predominately composed of CcmK2 may act as a barrier to CO_2_ leakage from the carboxysome into the cytosol, as well as O_2_ influx into the carboxysome, therefore facilitating the CO_2_-fixing activities and diminishing the oxygenation of Rubisco.

This mechanism of metabolite transit through the pore described in this study may be applicable to not only β-carboxysomes but also α-carboxysomes, based on the recent simulations study on the α-carboxysome shell protein CsoS1A and the β-carboxysome shell protein CcmK4^36^. Moreover, experimental studies have suggested that the α-carboxysome shell could impede the entrance of CO_2_ into the carboxysome^41^ and obviate its loss from the carboxysome lumen^43^.

Transit of the relatively large metabolites, such as 3-PGA and RuBP, through the pore of CcmK2 is potentially challenging. It remains to be tested whether it is possible for small molecules, or larger metabolites, or both, to move through the space between adjoining CcmK hexamers. Though the 3-PGA transit is energetically favorable (Fig. 3), it could induce a detectable conformational change of CcmK2 (Fig. 5). Given that RuBP is larger than 3-PGA, the RuBP transit through the CcmK2 pore is presumably more difficult. By contrast, the pseudo-hexamer CcmP, another shell protein in the β-carboxysome, bears a larger central pore (∼13 Å) compared with CcmK2^45,46^. It may act as a conduit for large metabolites, as a 3-PGA molecule may fit the pore of CcmP^46^ and a glycerol molecule has been observed in a binding pocket of the CcmP open structure^45^. The mechanism that controls RuBP and 3-PGA transit through the shell awaits further investigation. Moreover, cations, like Mg^2+^ that is required in the active site of Rubisco, may not be able to pass through the central pore of CcmK2 as easily as these large metabolites. It remains to be tested whether cations could transit through the CcmK2 pore or the pores of other shell proteins i.e. CcmK3, CcmK4 and CcmL, or through the gaps surrounding shell proteins generated by specific protein arrangement^34^ and dynamic self-assembly and interactions^25,47^.

Simulations revealed a 1-6 Å variation in the pore size of CcmK2 during metabolite passage (Fig. 5). This conformational change consists primarily of the loops forming the central pore region twisting, and the rotation of the Ser39 side chains into and away from the center of the pore. Greater variations in the pore size of shell proteins have been characterized in the open and closed forms of the pseudo-hexamers CcmP^45^, CsoS1D^48^, and EutL^49^. This suggests that pore conformational change could provide a means for selective metabolite transport (especially for large metabolite molecules) of shell proteins in the BMC counterparts. There was a distinct phenotype for each knockout mutant (Fig. 8, Supplementary Fig. 7). This may also suggest that there may be distinct roles for each shell protein or at least each class of BMC-T, BMC-P and BMC-H proteins, including the transit of the larger metabolites.

The rise of synthetic biology has spurred bioinspired engineering of BMC-based organelles for biotechnological applications, such as novel metabolic factories and molecule scaffolds, in light of the self-assembly, encapsulation and modular nature of BMC structures. Manipulating shell permeability is a pivotal strategy for design and construction of new BMCs to direct molecular transport and favor specific metabolic reactions. This study indicates that the mechanism controlling the permeability of the shell is complex and may be different between the different types of BMC as well as between the alpha and beta carboxysomes. The conserved residues of shell proteins that are vital for tuning the pore permeability, especially Ser39 of CcmK2, could be important targets in bioengineering to fine-tune molecular passage through beta-carboxysome shell structures. This study provides a predictive methodology to gain a more complete biophysical understanding of the mechanistic basis underlying molecular passage through the carboxysome shell. A deeper knowledge of the permeability of BMC shell proteins will inform strategies for the design and construction of new BMC-based nanoreactors to improve cell metabolism.

## Materials and Methods

### Cell culture and construction of genetic mutants

Syn7942 cultures were grown at 30°C under constant white light illumination of 30 μE·m^−2^·s^−1^, in BG11 medium^50^. Cultures were under one of three conditions, “Air” - atmospheric condition, “HCO_3_^−^” - BG11 supplied with NaHCO_3_ in every 24 hours to reach a final concentration of 150 mM, and “CO_2_” - 4% CO_2_ supplied in air. Cell growth was monitored at OD = 750 nm by a spectrophotometer (Jenway 6300). BG11 was supplemented with 10 mM TES buffer to increase its buffering capacity and minimize pH changes due to NaHCO_3_ addition^51^. *E. coli* cultures were grown at 37°C with constant shaking in Luria-Broth (LB).

Point mutation of CcmK2 (CcmK2-S39A) from Syn7942 was generated using the In-Fusion® approach in DH5α *E. coli*. Plasmids containing *ccmK2-S39A* and a kanamycin resistance cassette were transformed into Syn7942 cells to replace the native *ccmK2* gene in the genome, using Lambda RED Recombination^52^ as described previously^53–56^. Syn7942 knockout mutants Δ*ccmK3*, Δ*ccmK4*, Δ*ccmK3K4* and Δ*ccmP* were generated in a similar fashion by replacing the gene with a spectinomycin resistance cassette. Segregation of recombinant genes was verified by PCR and agarose gel electrophoresis. Mutants were selected for by the addition of 100 μg mL^−1^ kanamycin or spectinomycin where appropriate.

### Rubisco activity assays

*In vivo* carbon fixation from NaH^14^CO_3_ to ^14^C 3-PGA was monitored as previously described^11,56^. In brief, Syn7942 cells were incubated at 30°C with 25 mM NaH_14_CO_3_ for 2 minutes before permeabilization with 0.03 % (w/v) mixed alkyltrimethylammonium bromide (Sigma-Aldrich) and RuBP (Sigma-Aldrich) was then added to initiate the reaction. The reaction was terminated after 5 minutes and radioactivity measurements were carried out using a scintillation counter (Tri-Carb; Perkin-Elmer). Four biological repeats were prepared for each strain used in the kinetic analysis. Data processing was performed using SigmaPlot and figures were prepared using Origin.

### Crystal structure preparation

The full-length CcmK2 structures (PDB: 2A1B, 4OX7)^22,28^ and the CcmK2 C-terminal deletion structure (PDB: 3CIM)^27^ were obtained from the Protein Data Bank. Atoms in the unassigned regions of electron density maps of PDB structures were assigned manually and potential rotamers were fixed to a single conformation manually. Coot^57,58^ was used to view the electron density map and replace missing atoms and rotamers with our inferred most likely conformation.

### NAMD MD and US simulations

Umbrella sampling (US) molecular dynamics (MD) simulations were performed with the NAMD 2.12 program^59^ using the CHARMM36 force field (FF)^60^, to explore the passage of metabolites across the narrow central pore. Each system was built up using the CHARMM-GUI webserver^61^ and the ligands (HCO_3_^−^, CO_2_, O_2_ and 3-PGA) were automatically parameterized with CGenFF using their equilibrium structures obtained with the Gaussian 09 program package^62^ at the B3LYP/Def2-TZVP+GD3 level of theory. The protonation state of 3-PGA was chosen according to the microspecies distribution at pH = 7 predicted by the MarvinSketch software of ChemAxon. CcmK2-ΔC cyclic hexamer and the ligands were solvated in a hexagonal prism TIP3 water box of height 70 Å, then chloride ions were added to 0.15 M, and potassium ions were added to neutralize the system. For each system, corresponding to the four different types of metabolites, 10 molecules were randomly placed in the simulation box.

The MD protocol consisted of the following steps: (a) energy minimization over 2500 steps; (b) equilibration over 2 ns at constant pressure and temperature (*p* = 1.01325 bar, *T* = 303.15 K) with a root-mean-square deviation (RMSD) constraint of 1.0 kcal·mol^−1^·Å^−2^ applied on the protein backbone atoms; (c) 2 ns run in the NPT ensemble without constraints; (d) steered molecular dynamics (SMD) simulation - to obtain the starting positions for the subsequent umbrella sampling (US) simulations - with a pulling velocity of 10 Å ns^−1^, and SMD force of 7 kcal·mol^−1^·Å^−2^ applied on all heavy atoms of the small molecule; (e) 10 ns US production runs for each umbrella window in the NPT ensemble. Trajectories were run with a time step of 2 fs and the collective variables employed in US simulations were printed out in each step and used for the analysis. Constant temperature was set by a Langevin thermostat with a thermostat damping coefficient of 1 ps^-1^ with a collision frequency of 5 ps^−1^ ^62^. All of the bonds and angles involving hydrogen atoms were constrained by the SHAKE algorithm^62^. We used the particle mesh Ewald method^63^ for the long-range electrostatics in combination with a 12 Å cutoff for the evaluation of the non-bonded interactions.

The collective variable used as reaction coordinate (*z*) to describe the passage of ligands is defined as the projection of the molecule’s center of mass to the *z*-axis passing through the center of the pore (the positive *z* values correspond to the concave side of the protein). For each small molecule, 105 umbrella windows were run at steps of 0.5 Å (−26.0 ⍰ *z* ⍰ 26.0 Å) using a spring constant of 5.0 kcal·mol^−1^·Å^−2^. Moreover, the motion of the small molecule was constrained to the interior of a cylinder centered to the *z*-axis with *z*_min_ = −28 Å and *z*_max_ = 28 Å, whereas the maximum of the radial distance (*b*) from the z axis set to 15 Å. In case of Cl^−^ and K^+^ ions, 4 and 7 ions were selected in the cylinder, respectively by taking into account the *V*_cylinder_/*V*_simulation box_ ratio and concentration of ions. Their motion was completely free inside the cylinder (i.e. no umbrella bias applied) defined above and their trajectories interpreted as independent simulations during analysis. The free energy profiles and surfaces of substrate passage through the BMC pore, 1D(*z*) and 2D(*z*,*b*), were obtained using the weighted histogram analysis method (WHAM),^64,65^ and the dynamic histogram analysis method (DHAM),^65^ respectively. The mean and standard error for the 1D(*z*) free energy profiles were obtained by dividing the simulations for each US window into 5 equal-length segments, and analyzed these independently.

### Molecular dynamics and molecular docking simulations

Gromacs^66^ was used to perform 200 ns MDs of CcmK2-ΔC and CcmK2-full (PDB: 2A1B and 4OX7). All atom simulations were carried out in the GROMACS v2018.1 package with the CHARMM36 force field^60^ and the TIP3P model for the explicit solvent model. The leapfrog integrator was used to integrate the equations of motion every 2 fs. To restrict bond lengths the LINCS algorithm and PME with a real space cut-off at 14 Å were applied. RMSD was calculated by least square fitting of the protein to the crystal structure and then calculating the pairwise RMSD at each time step. RMSD was calculated for the entire hexamer assembly to take structural drift of the assembly in whole into consideration. Per residue RMSF were calculated for each chain and plotted as boxplots, to compare the flexibility of the three CcmK2 structures.

Two sets of independent docking simulations were carried out in SwissDock^38^ and Glide^39^, to characterize the binding poses of RuBP and 3-PGA with the 3CIM structure. Binding poses were scored using their FullFitness and clustered. Clusters were then ranked according to the average FullFitness of their elements.^67^

### Determining the pore diameter during metabolite flux

CPPTRAJ^68^ from Amber^69^ was used to align the trajectory files to the reference structure of CcmK2 (PDB: 3CIM) using the RMSD of alpha carbon atoms. The trajectories were then split into individual PDB files for each time frame. HOLE^37^ was used to measure the radius of the pore and given as an output of the diameter. This procedure was run for every timeframe that was split from the trajectory. These diameter measurements were made at the region corresponding to the minimum constriction along the pore axis *z*. VMD was used to find all the contacts, within a cutoff distance of 3.5 Å and angle of 30°, between each metabolite and the residues Arg11, Ser39 and Lys36. The number of contacts with each residue in each time frame was recorded. The pore diameter data was split into two categories, “no contact” when the metabolite did not fall within the cutoff of the specified residue, and “contact” when 1 or more contacts occurred. *P* values were calculated using the Wilcoxon rank-sum test from the SciPy python library, and violin plots were produced using Matplotlib.

### Autocorrelation analysis

To estimate the 2D-surface area of the pore, we defined a hexagon via the Ser39 CA atoms of the 6 loops defining the pore entrance that lie approximately on a plane (Supplementary Fig. 5A). We triangulated the hexagon via its vertices using a Delaunay Triangulation, and estimated the surface area as the sum of the areas of the triangles. We considered representative configurations from 3 US windows along the *z* axis when the metabolite is located at *z* = −20 Å, *z* = −5 Å, and *z* = 20 Å. The autocorrelation time, τ, associated with the temporal evolution of the pore surface area of CcmK2 from the US trajectories for the metabolites was measured for the selected US windows (Supplementary Fig. 5B).

### Protein sequence alignment

The protein sequences of shell hexamer homologs were taken from KEGG (https://www.genome.jp/kegg/) and the alignment was conducted using Clustal Omega.

### Computational resources

The computational work for this article was partially performed on resources of the National Supercomputing Centre, Singapore, the Chadwick Cluster, Advanced Research Computing, the University of Liverpool and ARCHER, the UK National Supercomputing Service.

## Acknowledgements

We thank Dr Fang Huang and Dr. Yaqi Sun for the technical assistance in generating knockout mutants. This work was supported by Royal Society (UF120411, URF\R\180030, RGF\EA\181061, RGF\EA\180233, and RG130442 to L.-N.L.), the Biotechnology and Biological Sciences Research Council Grant (BB/M024202/1 and BB/R003890/1 to L.-N.L.), the Engineering and Physical Sciences Research Council (EP/N020669/1 and EP/R013012/1 to E.R.), and the funding from A*STAR Graduate Academy (A*GA) Singapore.

## Author contributions

P.J.B., E.R., and L.-N.L conceived the project; M.F., I.S., S.L.W., F.S., and R.G.H. performed the simulations; M.F., I.S., S.L.W., F.S., and R.G.H. analyzed the data with input from P.J.B., E.R., L.-N.L.; M.F conducted the experimental studies and made the point mutations. M.F., S.L.W., E.R., and L.-N.L. wrote the paper with input from all authors.

## Additional Information

### Competing Interests

The authors declare no competing interests.

## Notes

### Competing Interest Statement

The authors have declared no competing interest.

