## Supplementary Information for "Molecular simulations unravel the molecular principles that mediate selective permeability of carboxysome shell protein"

**Supplementary Table 1. Carbon fixation activities of Syn7942 strains tested in this study**.

| **Strain** | ***Vmax*** **(μmol•min$^{-1}$•OD$^{-1}$)** | ***Km*****(RuBP) (mM)** |
|---|---|---|
| WT | 28.3 ± 2.0 | 0.7 ± 0.4 |
| Δ*ccmK3* | 17.4 ± 2.6 | 2.3 ± 0.7 |
| Δ*ccmK4* | 18.6 ± 1.6 | 2.3 ± 0.4 |
| Δ*ccmP* | 19.3 ± 11.1 | 7.7 ± 6.2 |
| Δ*ccmK3K4* | 3.6 ± 0.2 | 0.5 ± 0.1 |
| ΔccmK2-S39A | 9.9 ± 1.5 | 1.7 ± 0.6 |
| Δ*ccmK3*+CcmK2-S39A | 8.9 ± 1.6 | 1.4 ± 0.7 |
| Δ*ccmK4*+CcmK2-S39A | 4.8 ± 0.3 | 0.3 ± 0.1 |
| Δ*ccmP*+CcmK2-S39A | 2.9 ± 0.2 | 0.3 ± 0.1 |
| Δ*ccmK3K4*+CcmK2-S39A | 2.5 ± 0.4 | 0.7 ± 0.4 |

**Supplementary Table 2. Wilcoxon rank-sum test *P* values for metabolite-protein contacts.** The n value corresponds to the number of frames, when the substrate is either not within the contact cutoff of the residue, or when one or more atoms of the substrate is within the contact cutoff of the residue.

| **Substrate** | **Residue** | ***n* value** | | ***P*-value (Wilcoxon rank-sum test)** |
|---|---|---|---|---|
| | | **no contact** | **contact** | |
| $O_2$ | Arg11 | 2057 | 3 | 0.568 |
| | Lly36 | 2038 | 22 | 0.122 |
| | Ser39 | 2035 | 25 | 0.240 |
| $CO_2$ | Arg11 | 1185 | 5 | 0.890 |
| | Lys36 | 1184 | 6 | 0.277 |
| | Ser39 | 1167 | 23 | 0.086 |
| $HCO_3^-$ | Arg11 | 1958 | 422 | 0.698 |
| | Lys36 | 2111 | 269 | 0.0002 |
| | Ser39 | 1766 | 614 | 0.290 |
| 3-PGA | Arg11 | 1344 | 996 | $3.797e^{-13}$ |
| | Lys36 | 1856 | 484 | 0.013 |
| | Ser39 | 1241 | 1099 | $1.359e^{-17}$ |

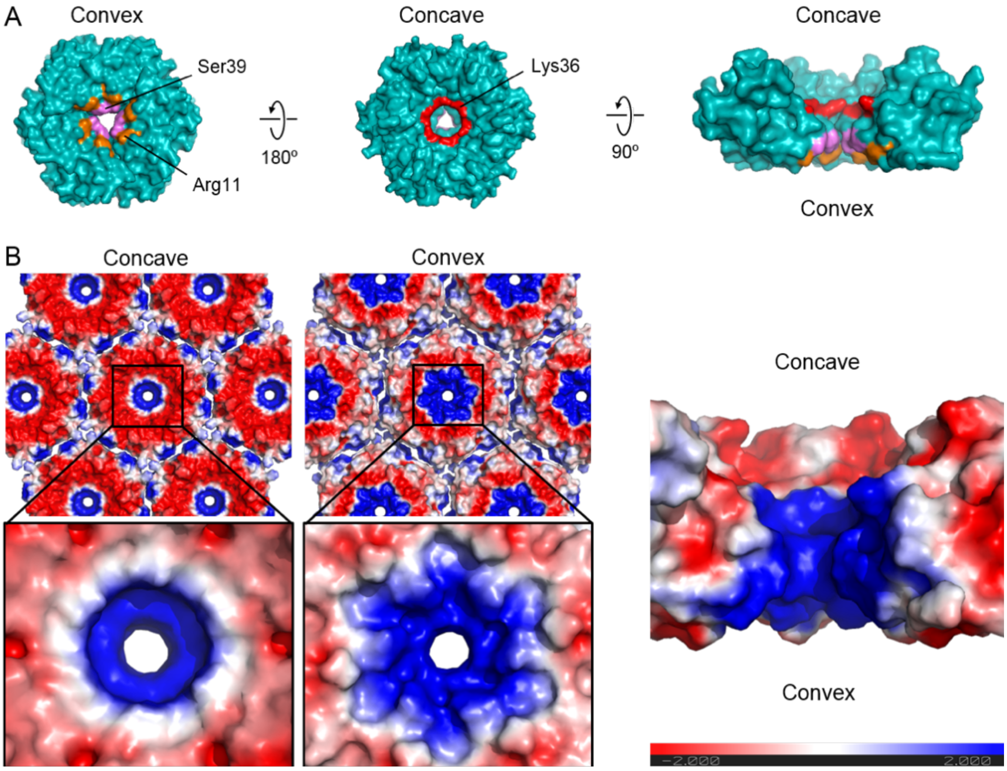


**Supplementary Fig. 1. Structural analysis of CcmK2.** A, Surface representations of CcmK2 (PDB: 3CIM) from different orientations, with Ser39, Lys36 and Arg11 residues highlighted in pink, red and orange, respectively. B, Electrostatic potential of CcmK2 (PDB: 2A1B), with the negatively charged residues in red and positively charged residues in blue. The concave side of CcmK2 is mostly negatively charged and a large area of the convex side has positive electrostatic potential, indicative of the charged-based tuning mechanism of the pore for molecular passage. The central pore is positively charged, due to the positively charged residues Lys36 and Arg11. The electrostatic potential was calculated using PyMOL with APBS plugin.

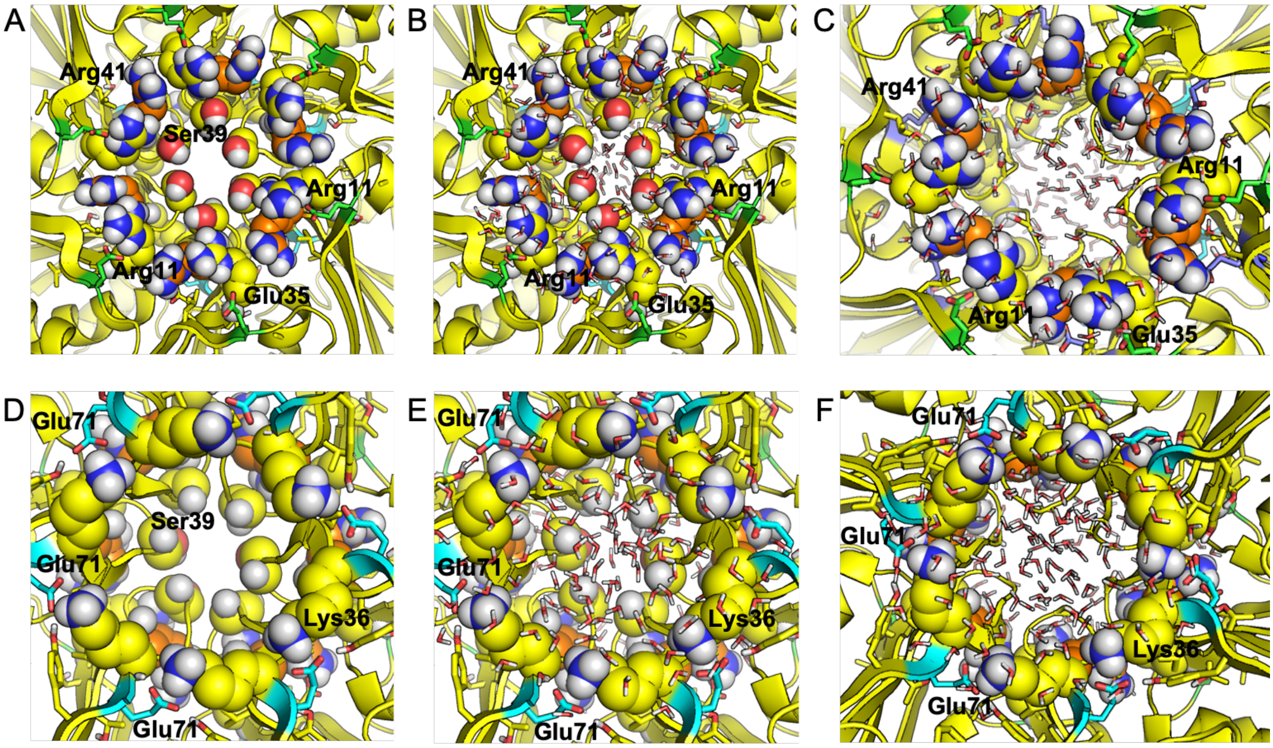


**Supplementary Fig. 2. Convex (A-C) and concave (D-F) sides of the pore**. Key positively charged Arg (A-C, yellow and orange VDW spheres) and Lys (D-F, yellow VDW spheres) residues are lined up at the pore entrance, directly stabilized by salt bridges via key Glu residues (green and blue sticks respectively) as well as solvent water molecules. Ser39 forms the bottleneck of the pore (A-B, D-E, yellow and red VDW spheres). Furthermore, water molecules also occupy the middle of the pore (B-C, E-F, sticks). Note that the convex side is lined up by two sets of positively charged residues (Arg11 and Arg41) occluding the pore entrance more than the concave side with Lys36 presenting a less charged environment and a larger diameter.

```
CcmK2-S.elongatus7942        ----MPIAVGMIETRGFPAVVEAADAMVKAARVTLVGYEKIGSGRVTVIVRGDVSEVQAS
CcmK2-Synechocystis6803      ----MSIAVGMIETRGFPAVVEAADSMVKAARVTLVGYEKIGSGRVTVIVRGDVSEVQAS
CcmK2-T.elongatus            ----MPIAVGMIETRGFPAVVEAADAMVKAARVTLVGYEKIGSGRVTVIVRGDVSEVQAS
CcmK1-Synechocystis6803      ----MSIAVGMIETLGFPAVVEAADSMVKAARVTLVGYEKIGSGRVTVIVRGDVSEVQAS
CcmK1-Synechocystis6714      ----------MIETLGFPAVVEAADSMVKAARVTLVGYEKIGSGRVTVIVRGDVSEVQAS
CcmK2-Synechocystis6714      --------------------------MVKAARVTLVGYEKIGSGRVTVIVRGDVSEVQAS
CcmK2-S.elongatus6301        ----MPIAVGMIETLGFPAVVEAADAMVKAARVTLVGYEKIGSGRVTVIVRGDVSEVQAS
CcmK3-S.elongatus7942        ----MPIAVGTIQTLGFPPIIAAADAMVKAARVTITQYGLAESAQFFVSVRGPVSEVETA
CcmK4-S.elongatus7942        ---MSQQAIGSLETKGFPPILAAADAMVKAGRITIVSYMRAGSARFAVNIRGDVSEVKTA
PduA-Salmonella              ---MQQEALGMVETKGLTAAIEAADAMVKSANVMLVGYEKIGSGLVTVIVRGDVGAVKAA
Hoch_5815-H.ochraceum        ----MADALGMIEVRGFVGMVEAADAMVKAAKVELIGYEKTGGGYVTAVVRGDVAAVKAA
CsoS1C-H.neapolitanus        MAAVTGIALGMIETRGLVPAIEAADAMTKAAEVRLVGRQFVGGGYVTVLVRGETGAVNAA
RmmH-M.smegmatis             ---MSSNAIGLIETKGYVAALAAADAMVKAANVTITDRQQVGDGLVAVIVTGEVGAVKAA
EutM-E.coli                  -----MEALGMIETRGLVALIEASDAMVKAARVKLVGVKQIGGGLCTAMVRGDVAACKAA

CcmK2-S.elongatus7942        VSAGLDSAKRVAGGEVLSHHIIARPHENLEYVLPIRYTEAVEQFRM----------
CcmK2-Synechocystis6803      VSAGIEAANRVNGGEVLSTHIIARPHENLEYVLPIRYTEEVEQFRTY---------
CcmK2-T.elongatus            VAAGVDSAKRVNGGEVLSTHIIARPHENLEYVLPIRYTEAVEQFRN----------
CcmK1-Synechocystis6803      VTAGIENIRRVNGGEVLSNHIIARPHENLEYVLPIRYTEAVEQFREIVNPSIIRR-
CcmK1-Synechocystis6714      VTAGIENIRRVNGGEVLSNHIIARPHENLEYVLPIRYTEAVEQFREIVNPSIIRR-
CcmK2-Synechocystis6714      VSAGIEAANRVNGGEVLSTHIIARPHENLEYVLPIRYTEEVEQFRTY---------
CcmK2-S.elongatus6301        VSAGLDSAKRVAGGEVLSHHIIARPHENLEYVLPIRYTEAVEQFRM----------
CcmK3-S.elongatus7942        VEAGLKAVAETEGAELINYIVIPNPQENVETVMPIDFTAESEPFRS----------
CcmK4-S.elongatus7942        MDAGIEAAKNTPGGTLETWVIIPRPHENVEAVFPIGFGPEVEQYRLSAEGTGSGRR
PduA-Salmonella              TDAGAAAARNV-GE-VKAVHVIPRPHTDVEKILPKGISQ-----------------
Hoch_5815-H.ochraceum        TEAGQRAAERV-GE-VVAVHVIPRPHVNVDAALPLGRTPGMDKSA-----------
CsoS1C-H.neapolitanus        VRAGADACERV-GDGLVAAHIIARVHSEVENILPKAPEA-----------------
RmmH-M.smegmatis             TEAGAETASQV-GE-LVSVHVIPRPHSELGAHFSVSSK------------------
EutM-E.coli                  TDAGAAAAQRI-GE-LVSVHVIPRPHGDLEEVFPIGLKGDSSNL------------
```

**Supplementary Fig. 3. Protein sequence alignment of BMC shell hexamer homologs**. The three key residues of interest Ser39, Lys36, and Arg11 are highlighted in pink, red, and orange, respectively. Ser39 and Lys36 in the pore motif of K-I-G-S are conserved among shell protein homologs. Arg11 is located at the pore entrance on the convex side of CcmK2 along with Arg41 that is highlighted in aqua.

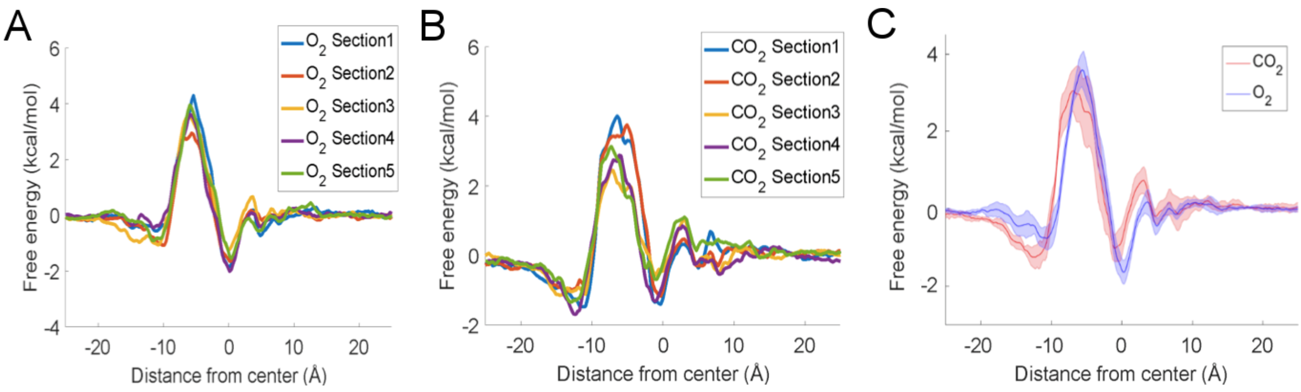


**Supplementary Fig. 4. Free energy profile reconstruction**. To assess the convergence of the US simulations in the reconstruction of the representative free energy profiles, we divided each US simulation window into 5 equal-length trajectories (sections) and analyzed them independently, as shown for the $O_2$ molecule (A) and the $CO_2$ molecule (B). We checked the independent profiles did not display significant systematic changes and calculated the mean and standard error for the representative 1D($z$) free energy profiles (C).

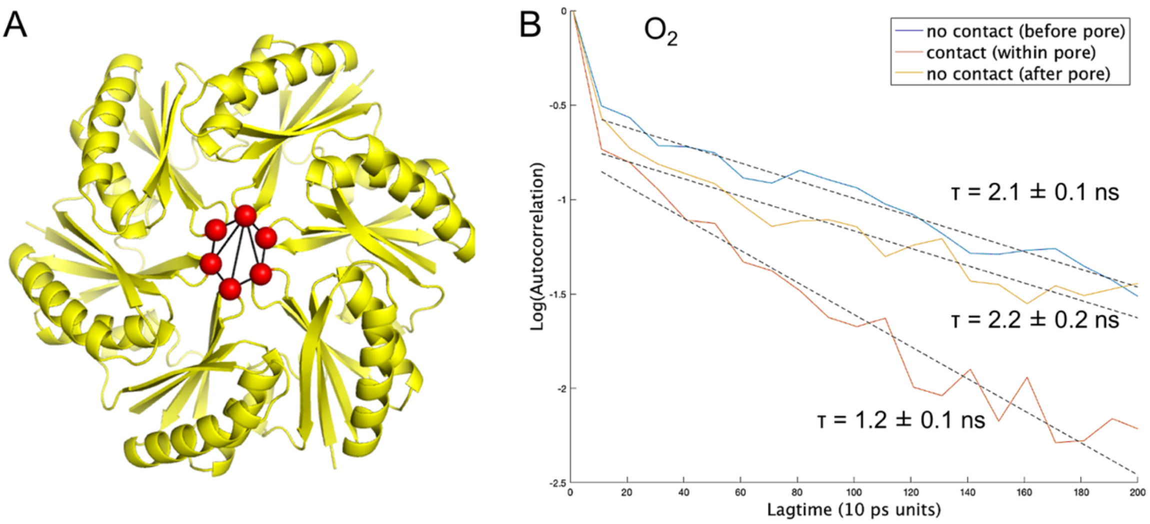


**Supplementary Fig. 5. Delaunay triangulation of the pore and autocorrelation.** A, Triangulated hexagon defined via the Ser39 CA atoms of the 6 loops (red points) defining the pore entrance that lie approximately on a plane, used to estimate the 2D surface area of the pore. B, The autocorrelation time, $\tau$, associated with the temporal evolution of the pore surface area of CcmK2 from the Umbrella Sampling trajectories for $O_2$. We considered 3 representative US windows along the Z axis when the metabolite is located at Z = -20 Å (blue, convex side), Z=-5 Å (red, within the pore region), and Z = 20 Å (yellow, concave side). We expect that no large conformational changes take place orthogonal to our collective variable. As the pocket size is relatively large, the US windows are long enough to enable the smaller conformational changes during our MD simulations either driven by the US reaction coordinate, or, if orthogonal to this, during the MD simulations in each window. Nevertheless, we cannot exclude that at even longer timescales further conformational changes may occur at the pore, especially taking into account other components of the system (e.g., C-terminus, or other proteins interacting with the shell) not included in our simulations.

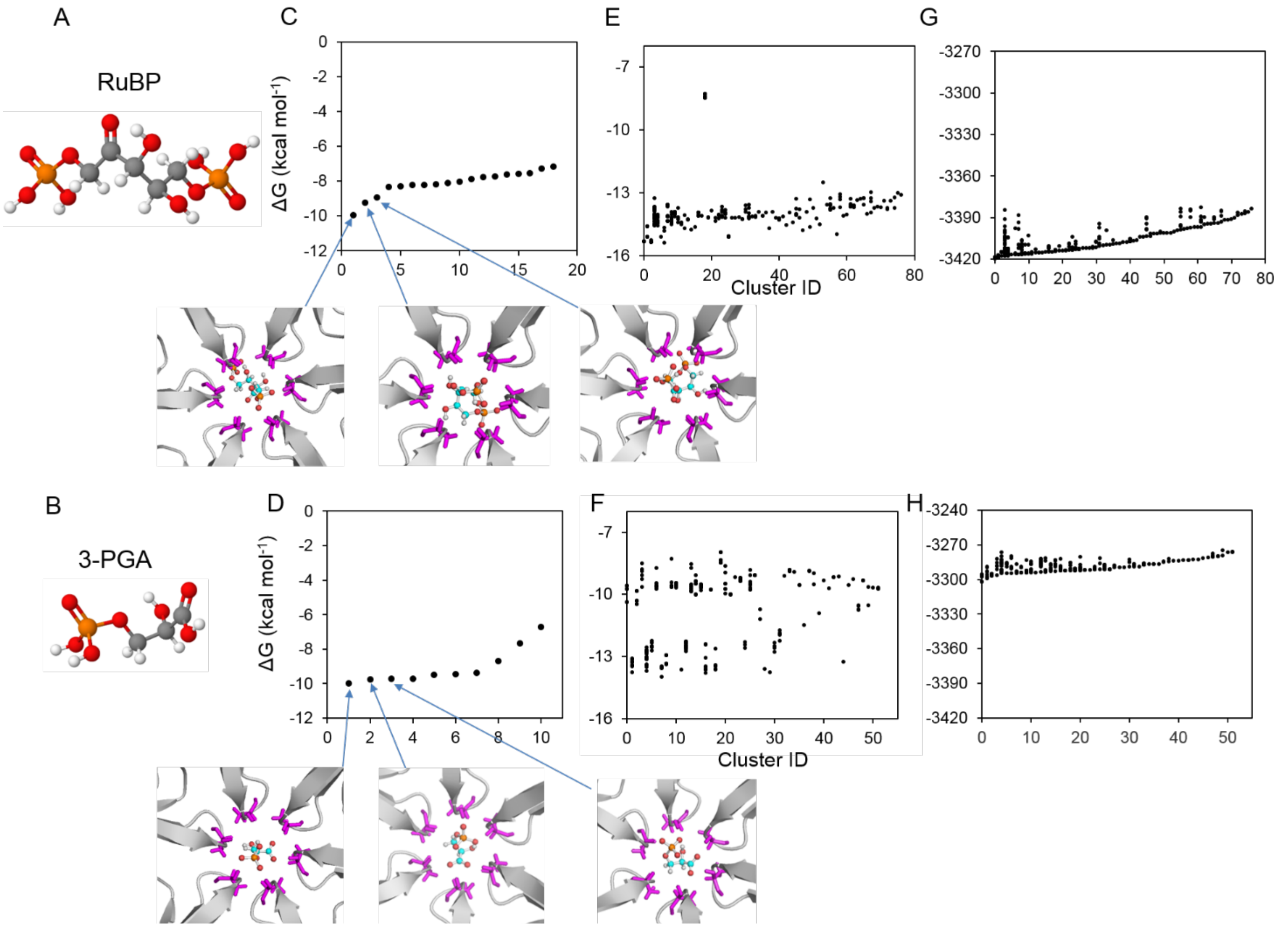


**Supplementary Fig. 6. Molecular docking of RuBP and 3-PGA with CcmK2 using SwissDock and Glide SP**. A-B, Molecular structures of RuBP and 3-PGA. C-D, Free energy profiles of RuBP and 3-PGA docking to CcmK2 using Glide XP, the three lowest free energy poses are shown underneath, and the corresponding data points are indicated by blue arrows. E-F, Free energy profiles of RuBP and 3-PGA docking to CcmK2 using SwissDock. G-H, FullFitness plots of RuBP and 3-PGA using SwissDock.

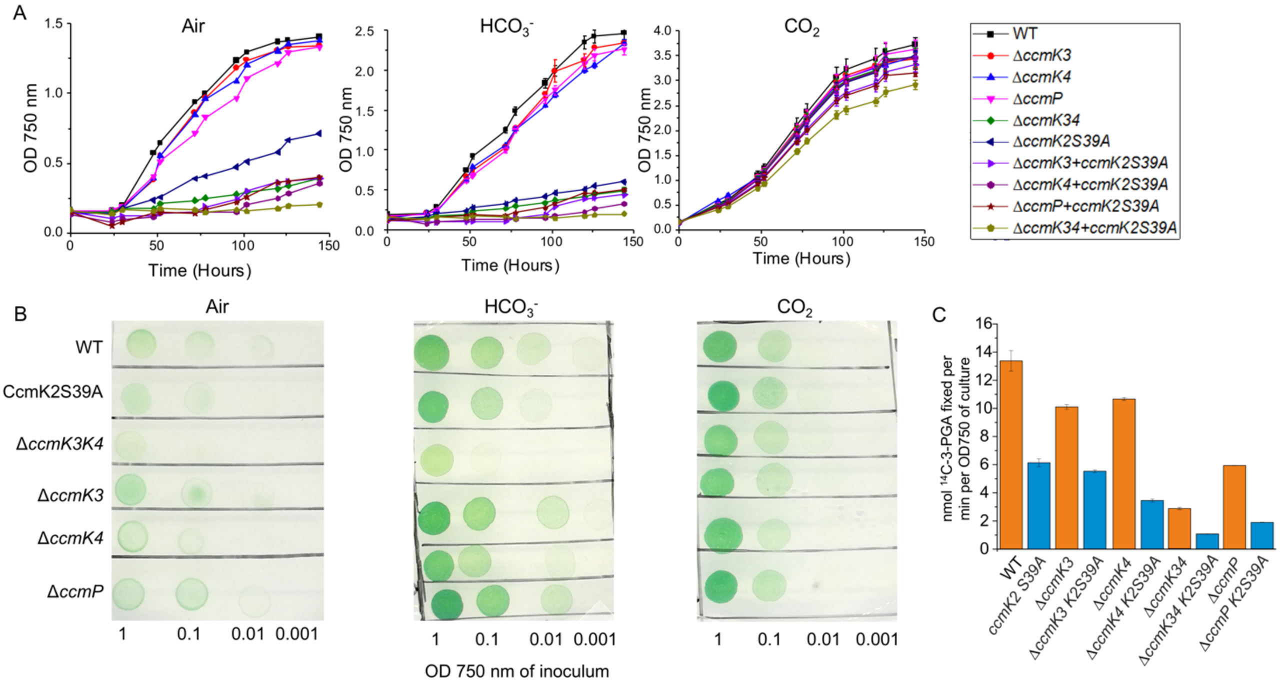


**Supplementary Fig. 7. Physiology of strains in air, $HCO_3^-$ and $CO_2$ conditions**. A, Growth curves of strains in air, $HCO_3^-$ and 4% $CO_2$ conditions, respectively ($n$ = 3). B, Spot assays of the strains grown on BG11 plates in air, $HCO_3^-$ and 4% $CO_2$ conditions ($n$ = 2) after 48 hours. C, Rubisco activities of cells grown under 4% $CO_2$, measured by $^{14}C$ radiometric carbon fixation assay at 0.4 mM RuBP ($n$ = 3).
